# Dynamics of cell mass and size control in multicellular systems and the human body

**DOI:** 10.1101/2020.12.03.411017

**Authors:** David Martinez-Martin

## Abstract

Cellular processes, in particular homeostasis and growth, require an intricate and complex exchange of matter between a cell and its surroundings. Yet experimental difficulties have prevented a detailed description of the dynamics of a cell’s mass and volume along different cellular processes, limiting our understanding of cell physiology in health and disease. It has been recently observed that single mammalian cells fluctuate their mass in a timescale of seconds. This result challenges central and long-standing cell growth models, according to which cells increase their mass either linearly or exponentially throughout the cell cycle. However, it remains unclear to what extent cell mass fluctuations may be sustained in multicellular organisms. Here I provide a mathematical model for cell mass fluctuations and explore how such fluctuations can be successfully sustained in multicellular organisms. I postulate that cells do not synchronise their mass fluctuations, but they are executed with their phases uniformly distributed. I derive a mathematical expression to estimate the resulting mass shift between fluid compartments in an organism due to cell mass fluctuations. Together with a new estimate of 4×10^13^ human cells in the body, I demonstrate that my hypothesis leads to shifts of mass between the intracellular and extracellular fluid compartments in the human body that are approximately or smaller than 0.25 mg and, therefore, perfectly viable. The proposed model connects cell physiology with information theory and entropy.

## Main text

Over the last two centuries it has become well accepted that cells are the most basic units of life (1) and, consequently, that all known living organisms are made up of cells. During this time, important scientific efforts have led to breakthroughs about the composition, structure and functioning of cells and their components (2). More recently, the strong development of omic technologies with single-cell resolution makes it possible to accurately characterise complex processes such as gene-expression dynamics, cell heterogeneity or lineage tracing (3-5).

Nevertheless one of the most fundamental processes of life, cell growth, which I here describe as the dynamics and regulation of a cell’s mass along different biological processes, remains largely unclear (6), limiting our understanding of the development and functioning of single and multicellular organisms. Importantly, dysregulation of cell mass is linked to aging and major health conditions including cancer, diabetes, cardiovascular diseases, and obesity (7-12). Accurately identifying cell mass dynamics is needed to advance our understanding of health and disease, and ultimately to improve diagnosis and treatment for people with health conditions.

The density of a cell, which corresponds to the total mass of a cell divided by the cell’s total volume, is close to that of water and it is subject to small variations (13), generally smaller than 5-6% as recorded in yeast (12). Specially, the density of mammalian cells, and in particular human cells, is largely constant over the cell cycle with the highest variation of approximately 0.5% during mitosis (13). As example, the density of murine lymphocytic leukaemia cells remains between 1.060 g/cm^3^ and 1.070 g/cm^3^ over the entire cell cycle, including mitosis (14). Notably, one must be careful when interpreting the literature, as sometimes such variations are reported subtracting the density of water (1.000 g/cm^3^) from the cell density values, which make the numbers look larger (12), or the term density is used to designate other magnitudes. Considering the fact that the density of mammalian cells is largely constant, changes of a mammalian cell’s total mass translate with an accuracy higher than 1% into changes of cell volume, the latter being the historically preferred magnitude because it can be more readily assessed using common technologies including optical microscopies, coulter devices and flow cytometers. Although such technologies can provide valuable measurements of a cell’s volume, they cannot continuously track volume changes in real time (millisecond time resolution) and with high accuracy (<1% of total cell volume), limiting our description of the dynamics and regulation of cell volume and mass.

Until now, two mathematical models are mainly used to describe the dynamics of a cell’s mass and volume over the cell cycle: a linear and an exponential model (Fig 1a). The linear model implies that cells increase their mass at a growth rate (derivative of mass with respect time) that is constant over the cell cycle, whereas the exponential model implies that the rate of mass change is proportional to the cell mass at any given time. These models are protagonists of a long lasting and active controversy (6, 9, 15-20), which lies on the theoretical basis that a linear model of cell growth does not require the cells to have an active regulation mechanism of cell mass or volume whereas the exponential model does (6, 21). Interestingly, both models imply a monotonic behaviour of a cell’s mass over time, and exclude the possibility of intervals with negative growth in which a cell can decrease its mass (and volume) at a given time (Fig 1b). Cell growth monotonicity has been universally accepted and so, within the context of single cells, cell growth is defined as mass accumulation (8).

**Figure 1.**
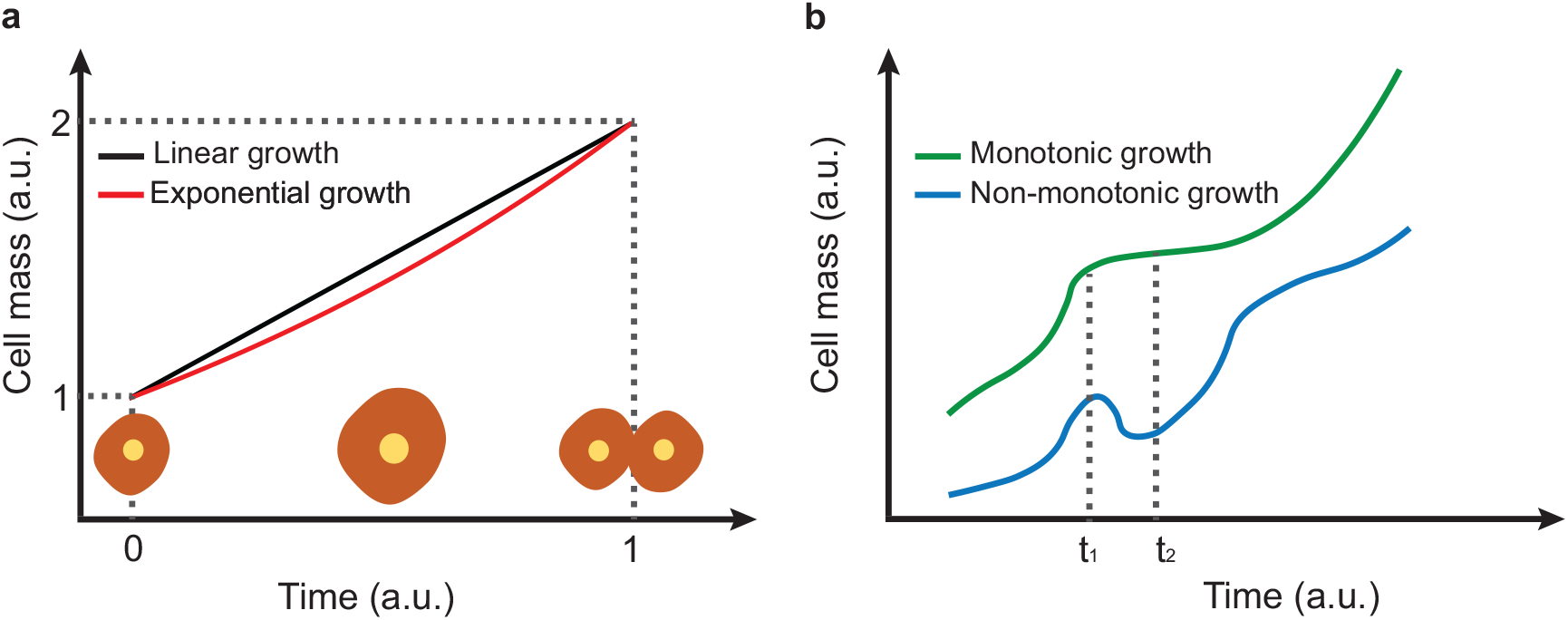
Cell mass and growth. **a**, Representation of a cell changing its mass throughout the cell cycle following traditional models. The black curve shows a linear growth and the red curve shows an exponential growth. The cell eventually divides into two daughter cells. **b**, Examples of monotonic and non-monotonic mass growth behaviours. A monotonic increasing (decreasing) behaviour means that a cell’s mass at any given time, t_1,_ is lower (higher) or equal than the cell’s mass at any other time, t_2_, where t_1_≤t_2_. Both central models in (a) imply a monotonic mass increase. Recently identified mass fluctuations (25) in mammalian cells challenge monotonicity.

A recently developed inertial picobalance technology (22-24) allows continuously measuring the total mass of single and multiple cells with picogram accuracy and millisecond time resolution. Using this technology it has been shown that despite mammalian cells develop an overall increase of mass throughout the interphase in the time scale of hours, such cells fluctuate their total mass by approximately 15 pg in a much shorter time scale of a few seconds (25), challenging the central cell growth assumption of monotonic behaviour for cell mass and volume. Initial perturbation experiments indicate that these mass fluctuations, which remain also during cell growth arrest induced by Vaccina virus infection, are actively driven and largely mediated by water transfer in and out of the cells (25). This makes it very difficult to detect such mass fluctuations with other cell mass sensing technologies, particularly because they are much more limited in time resolution, and/or total mass resolution (25). Moreover, they generally do not directly measure a cell’s total mass but a small fraction of it, such as the buoyant mass or the dry mass (26). It is important to note that the buoyant mass corresponds to the difference between a cell’s total mass and that of the volume of media displaced by the cell, and is therefore in the order of only 5% of a cell’s total mass (27). The dry mass corresponds to the total mass of a cell minus the mass of its water content, corresponding to approximately 30% of a cell’s total mass (13).

The newly identified fluctuations of a cell’s total mass (25) provides direct and robust experimental evidence that cells actively regulate their own mass over time, and their study is key to better understand cell physiology. These fluctuations may be directly intended or the consequence of a regulated process (28, 29), which suggests that cells control their mass by at least a negative feedback, which may have an adaptive set-point involved in the regulation of cellular functions. Indeed, changes of the intracellular water content and hence mass modulate the concentration and interaction of intracellular molecules and structures, and are linked to the regulation of cellular processes including metabolism, gene expression, cell proliferation or cell death (30-32). Additionally, fluctuations of a cell’s mass may be the result of multiple feedbacks or time delays (28, 29). Besides, cells may employ mass fluctuations to interact with the environment and/or sense their own mass. Yet, the discovery of single cell mass fluctuations raises a fundamental question: To what extent such fluctuations may be sustained in multicellular organisms including the human body?

Guided by Shannon’s information theory (33) and Jaynes’ principle of maximum entropy (34, 35), I propose that cells do not fluctuate their mass synchronously, but with the phases of their fluctuations following a uniform probability distribution. Jaynes’ principle establishes that the best representation of our knowledge about the state of a system is given by the probability distribution with the highest entropy subject to the existing knowledge (35). Currently we do not know constrains for the phases of cells’ mass fluctuations and the highest entropy corresponds to a uniform probability distribution (36). Given my hypothesis, I derive a mathematical inequality to estimate the mass shift between the intra- and extracellular fluid compartments in multicellular systems. Moreover, I provide a new estimate for the number of human cells in the body and mathematically demonstrate that cell mass fluctuations can be sustained in the body with negligible resulting shifts of mass between body fluid compartments. Thus mass fluctuations are a perfectly viable process that preserves homeostasis whilst enabling the interaction and communication of cells with their environment.

Considering that mammalian cells fluctuate rapidly their mass with certain periods or frequencies (25), the fluctuated mass can be described using the formalism of harmonic oscillators. For simplicity and without loss of generality, I assume that cells fluctuate mass at one single frequency (Fig. 2) within a given time frame. However, the same formalism can be applied if cells fluctuate mass at multiple frequencies.

**Figure 2.**
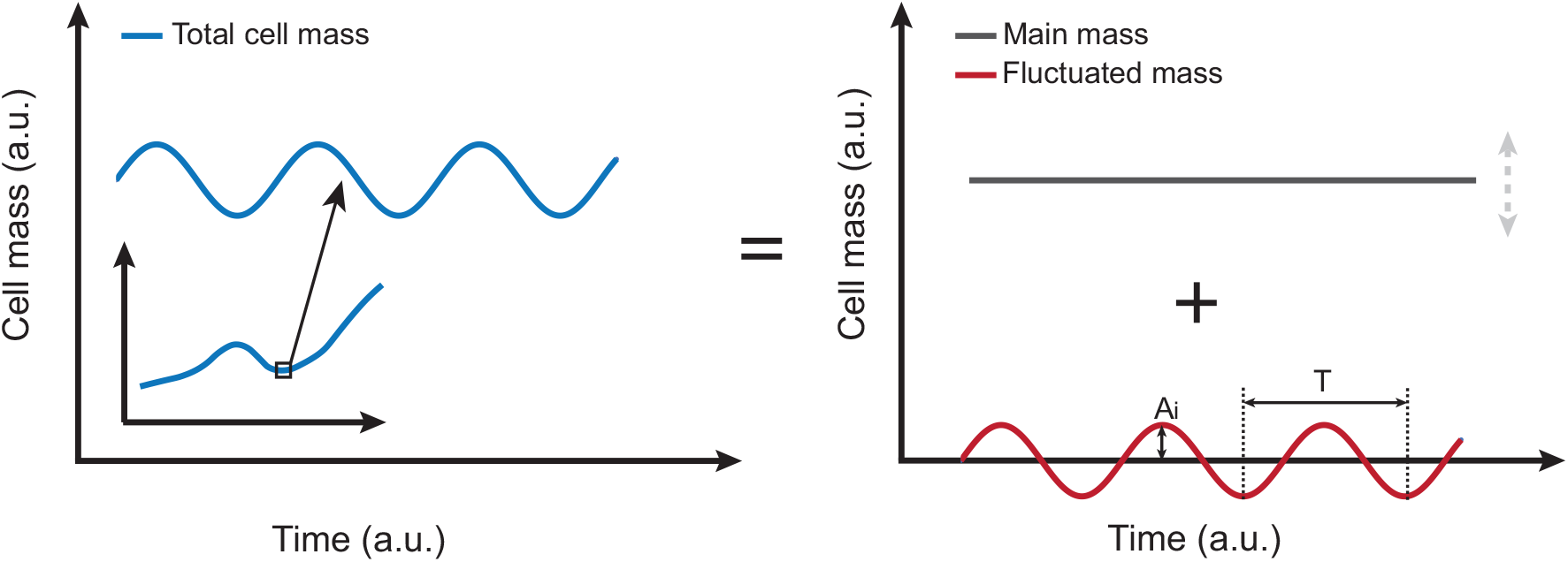
Fast fluctuations of a cell’s mass. The inset (left plot) illustrates the dynamics of a cell’s total mass over an extended period of time (∼hours). The left plot corresponds to a zoom in of the previous inset in which fluctuations of a cell’s mass are visible over a span of tens of seconds. As depicted in the right plot, the fluctuations can be decomposed into two components: the fluctuate mass, which shows certain periodicity (T) and amplitude (Ai), and the main mass. The latter may also vary over time as shown by the grey arrow (right plot), yet at much slower rate than the fast fluctuated mass (timescale of seconds). Thus it can be considered largely constant for the effects analysed here.

Taking into account the previous considerations, let *ψ*_*i*_ be the fluctuated mass of a certain cell *i* within a given window of time; *A*_*i*_ is the amplitude of the mass fluctuations; *ω* corresponds to the angular frequency of the mass fluctuations, which is equal to 2*π*/*T*, being *T* the period of the fluctuations; *t* represents time; and *δ*_*i*_ corresponds to its phase. We can then write

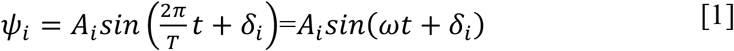

Mammalian cells forming an organism are surrounded by interstitial fluid, which comprises the extracellular fluid virtual compartment. Additionally, the total amount of water within all cells constitutes the intracellular fluid virtual compartment. Fast cell mass fluctuations involve water exchange in and out of the cells and must preserve the total mass of the organism they are part of. Therefore, the mass fluctuations of extracellular fluid (*χ*) as a result of cell mass fluctuations plus the mass fluctuations of all the cells in the organism must equal zero, with *N* being the number of mammalian cells in a given organism:

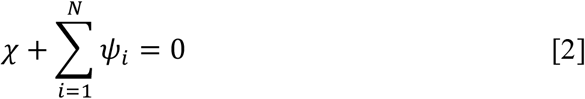

The second term in Equation [2] is a sum of sinusoidal functions with the same frequency. Such term can be written as another sinusoidal function based on the harmonic addition theorem.

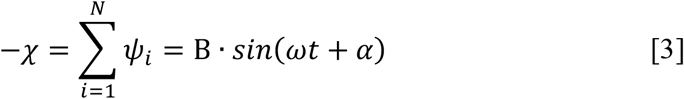

where *B* is the amplitude of the mass fluctuations of the intracellular fluid compartment and *α* the phase of such fluctuations. Making use again of the harmonic addition theorem (37), the square of the amplitude of the intracellular fluid fluctuations can be written as

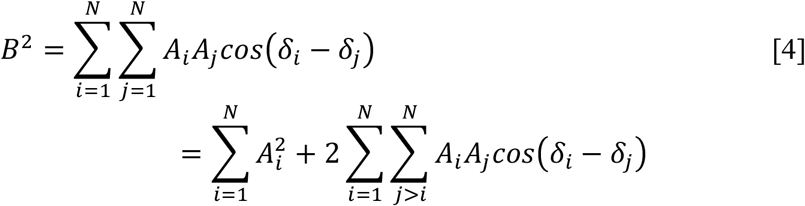

Importantly, given the hypothesis I propose in this article, the phases of the fluctuated mass of cells are given by a circular uniform probability distribution in the range [0, 2*π*). The amplitudes of the fluctuations are expected to be similar at least for cells of similar types (25). Hence for organisms comprising large numbers of cells of the different cell types forming the organism, the last term of the previous equation vanishes since the average value of the cosine is 0. Consequently,

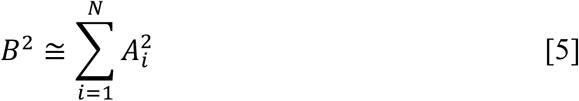

Let be *a* the highest amplitude of mass fluctuations among all the cells in the organism. That means that any cell will fluctuate its mass with an amplitude equal or smaller than *a*, i.e.

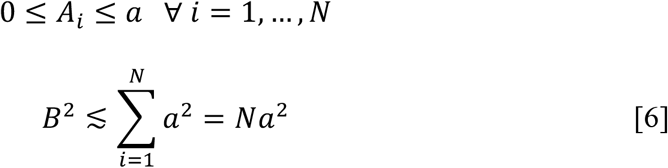

And taking the square root of each term I reach the final result,

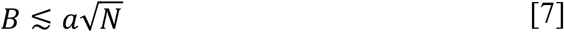

This powerful inequality estimates the amplitude of the intracellular fluid compartment fluctuations (*B*) as a consequence of cell mass fluctuations. Given equation [2], the same estimate is valid for the amplitude of the extracellular fluid compartment fluctuations.

Using Equation [7] one can estimate *B* in the particular case of a human being, which requires an estimate of the number of cells in the human body. The average mass of a human body is 70 kg (38), from which 60% is water (39), implying 42 kg of water. About 2/3 of the total amount of water corresponds to intracellular water and 1/3 is extracellular water (39). That means 28 kg of water are distributed within the cells and 14 kg is extracellular water. The average mass of a human cell is 1 ng (38), from which ∼70% is water (13). Consequently, the average amount of intracellular water per cell is 0.7 ng. We can then estimate the number of cells in the human body by dividing the total mass of intracellular water in a human being by the average mass of intracellular water per cell, which leads to *N* = 4 · 10^13^cells, and this is in very good agreement with recent estimations (38, 40). As recently identified by Gabriel S. Adams et al. people tend to overlook new solutions with advantageously subtractive changes (41), missing the opportunity to make science more simple, translatable and applicable when possible. Thus, an important feature of this new estimate is its genuine simplicity.

If every human cell in the body fluctuates its mass with an amplitude equal or smaller than *a* = 40 *pg* = 40 · 10^−12^*g*, which is a reasonable higher end amount (25) and approximately 4% of the total mass of an average human cell, the resulting fluctuations of the intra- and so extracellular fluid compartments have an amplitude that is approximately or smaller than 0.25 mg. Such amount is the aftermath that at any given time some cells uptake mass while others release it lowering the resulting mass shift. This amount is negligible compare to both the 28 kg of the intracellular fluid compartment and the 14 kg of the extracellular fluid compartment, making it a perfectly viable process.

Single cell mass fluctuations have been detected in mammalian cells in culture, and according to the model presented here, it is expected to be challenging to detect them directly by looking at the resulting collective behaviour of systems with large numbers of cells. I am hopeful that this work will stimulate the advancement and development of experimental techniques to be able to track in real-time the mass of single cells in vivo. It is important to note that fluctuations of a cell’s mass bring new possibilities to better understand molecular crowding, cell physiology, cell communication and developmental biology. Such quick exchanges of matter, which imply variations of approximately 1-4% of a cell’s mass within a few seconds that are largely comprised of water, are expected to have a dynamic impact in molecular crowding and concentration of chemical cellular components (42, 43), whose variations can be higher than 1-4% if such water-based fluctuations are not homogeneously distributed within the cell (44). Hence, such fluctuations can be involved in phase separation events (45, 46) and regulation of cellular functions (32). Moreover, due to the highly constant density of mammalian cells (13), said mass fluctuations involve similar cell volume fluctuations that may act as biomechanical signals in cell-cell communication (47). Therefore, such processes may be key to understand how cells signal and recognise their location within multicellular organisms. Thus cell mass fluctuations may be an effective way for cells to sample the environment but also to release signals, ensuring cell communication across cells forming an organism without compromising viability and homeostasis of the organism itself. For this to happen it is expected that cells fluctuate their masses asynchronously from each other. Yet, there may be circumstances in which the phases of such fluctuations may deviate from a uniform distribution and hamper homeostasis. Consequently, a better understanding of single-cell growth, which I define here as real-time single-cell mass, present a strong potential to be a universal biomarker bringing opportunities to improve our understanding of health and disease and ultimately to develop more accurate diagnostics and better treatments.

## Conclusion

I conclude that the model I propose here establishes the possibility for cells to be able to fluctuate and exchange mass rapidly with their surroundings, which may enable them to sense and interact with the environment, drive and regulate cellular processes, and detect their own mass. Moreover, I anticipate that cells fluctuate their mass with their phases uniformly distributed. I have demonstrated that in multicellular systems with large number of cells, such behaviour enables cells to exchange 1-4% of their total mass in the time scale of seconds while producing very small resulting mass shifts between fluid compartments, hence preserving homeostasis. Loss of such condition might contribute to excessive mass shifts between the intracellular and extracellular fluid compartments, which is known to be linked to health conditions (31, 48). I believe that this model, which connects information theory and statistical mechanics with cell physiology, brings new insight together with a powerful and unprecedented perspective about the dynamics and control of cell mass and volume. I envision that this work will promote further research and will stimulate a paradigm shift in multiple areas including cell physiology and developmental biology.

## Acknowledgments

This work has been supported by the SOAR award of the University of Sydney. I thank Natasha Tomm and Allison Tong for carefully reading the manuscript.

## Author contributions

D.M.-M conceived and developed the model, and wrote the manuscript.

## Ethics approval and consent to participate

N/A

## Consent for publication

The author provides consent for publication

## Availability of data and material

N/A. This is a theoretical paper

